# MutaFrame - an interpretative visualization framework for deleteriousness prediction of missense variants in the human exome

**DOI:** 10.1101/2021.06.03.446874

**Authors:** François Ancien, Fabrizio Pucci, Wim Vranken, Marianne Rooman

## Abstract

**Motivation:** High-throughput experiments are generating ever increasing amounts of various -omics data, so shedding new light on the link between human disorders, their genetic causes, and the related impact on protein behavior and structure. While numerous bioinformatics tools now exist that predict which variants in the human exome cause diseases, few tools predict the reasons why they might do so. Yet, understanding the impact of variants at the molecular level is a prerequisite for the rational development of targeted drugs or personalized therapies.

**Results:** We present the updated MutaFrame webserver, which aims to meet this need. It offers two deleteriousness prediction softwares, DEOGEN2 and SNPMuSiC, and is designed for bioinformaticians and medical researchers who want to gain insights into the origins of monogenic diseases. It contains information at two levels for each human protein: its amino acid sequence and its 3-dimensional structure; we used the experimental structures whenever available, and modeled structures otherwise. MutaFrame also includes higher-level information, such as protein essentiality and protein-protein interactions. It has a user-friendly interface for the interpretation of results and a convenient visualization system for protein structures, in which the variant positions introduced by the user and other structural information are shown. In this way, MutaFrame aids our understanding of the pathogenic processes caused by single-site mutations and their molecular and contextual interpretation.

**Availability:** Mutaframe webserver at http://mutaframe.com

---

Whereas the amount of genetic data obtained through high-throughput sequencing experiments has exploded in the last twenty years [1], it remains challenging to accurately predict and interpret how some gene variants lead to diseases, which are often caused by changes in the protein(s) the gene encodes [2]. Especially difficult to predict are the changes these variants cause at the level of protein behavior, which can often explain the pathogenic mechanisms involved and allows optimizing the rational development of targeted drugs. Multiple bioinformatics tools have been developed to classify variants in the human exome as deleterious or neutral [3, 4], but their explanatory power remains limited.

We present a substantial extension of the Mutaframe webserver [5], which is designed to improve the interpretability of such protein-level predictions via an easy-to-use graphical interface (Fig. 1). The new version features two complementary state-of-the-art predictors, DEOGEN2 [5, 6] and SNPMuSiC [7]. DEOGEN2 is a protein sequence-based predictor that utilizes evolutionary information as well as contextual information, such as the relevance of the gene containing the variant or the interactions of the encoded protein. SNPMuSiC uses as input experimental or modeled 3-dimensional (3D) protein structures and predicts deleterious variants on the basis of the changes in stability these cause.

**Figure 1:**
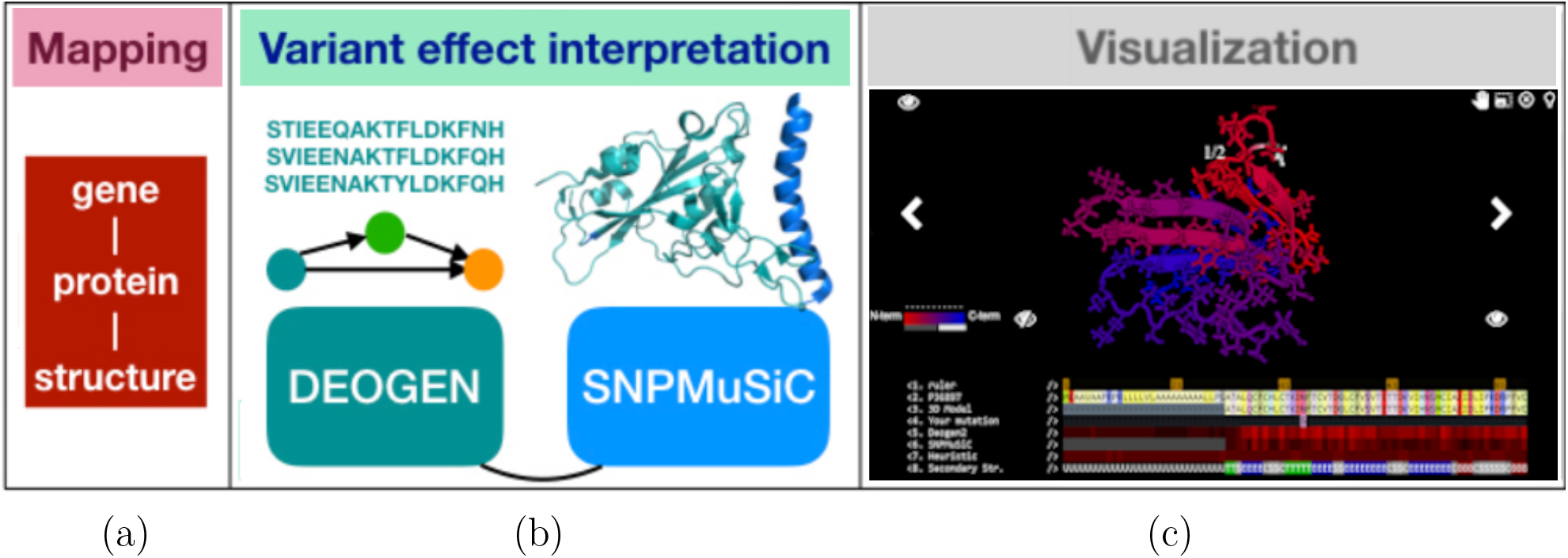
The capabilities of the MutaFrame webserver at the mapping (a), variant effect interpretation (b) and 3D structure visualization (c) levels (see also Supplementary Material).

Combining these two predictors, which already individually have good performances [3, 4], yields a consensus predictor with a balanced accuracy of 92% and a positive predictive value of 97% on 80% of the variants. Moreover, the combination of the explanatory power of DEOGEN2 in terms of evolutionary and contextual features, and of SNPMuSiC in terms of structure and stability, improves the contextualisation of the impact that a mutation has at the protein level. For example, highly conserved residues located in the protein core, whose variants are predicted as deleterious by both DEOGEN2 and SNPMuSiC, are highly likely to be destabilizing, thus inducing (partial) unfolding of the protein. A full description of these predictors, their performance, large-scale applications and case studies related to the Niemann-Pick disease is available from Supplementary Material.

The new version of the MutaFrame server also provides additional computational and visualization utilities that help the users in the interpretation of the prediction results:

- Visualization of the experimental or modeled 3D structure of the wild-type target protein, if available, and of the localization of the variant residue.
- Per-residue solvent accessibility and secondary structure as well as additional information on the 3D protein structures such as the resolution of the X-ray structure or of the template used for the homology modeling.
- DEOGEN2 and SNPMuSiC prediction scores of specific variants introduced by the user.
- Heatmap showing the DEOGEN2 and SNPMuSiC scores of all possible variants in a target protein, both along the sequence and in the 3D protein structure.
- Influence of the different features (residue conservation, protein essentiality, ..) in the DEOGEN2 prediction.
- Mapping between gene, protein sequence and protein structure identifiers, and corresponding sequence alignments, for the entire human proteome.

Note, moreover, that all the results available on the webserver can easily be downloaded for offline analyses. In summary, MutaFrame facilitates the analysis of human variants at the molecular, evolutionary and contextual levels, thus going beyond the simple binary deleterious/benign classification. This constitutes an important asset in the clinical and biopharmaceutical fields.

## Supplementary Information

Although major advances have been made over the past two decades in predicting whether missense variants in the human exome are deleterious or benign, it remains challenging to interpret their molecular impact and understand their role in pathogenic mechanisms.

In this paper, we present the MutaFrame web server which is designed to help improve our understanding of the effects of variants at the molecular level by providing a series of variant **Mapping, Interpretation** and **Visualization** utilities. The integration of these tools in the MutaFrame framework and its easy-to-use structure are two main characteristics that make it a powerful instrument for variant interpretation. A summarized description of these three utilities is given below:

- **Interpretation**. This utility is built around the results of two complementary predictors, DEOGEN2 [5] and SNPMuSiC [7], which both predict the deleteriousness of human exome variants and their impact on the disease phenotype. In addition, this utility provides information related to protein sequence and structure and to biophysical and contextual features that drive the prediction, so giving insights into why variants are predicted to be deleterious or neutral. This information is invaluable to better understand the impact the mutations might have.
- **Mapping**. Due to the increasing amount of biological data from *e.g*. genomics, proteomics and transcriptomics studies, it has become essential to link and cross-reference the different sources of data and annotations, and integrate them into a common framework. More specifically, this utility maps variants at multiple levels, *i.e*. in gene sequences, protein sequences and protein structures. Thus, even though MutaFrame primarily focuses on proteins, this utility retrieves information about variant characteristics across several scales, which again can help to contextualise and explain the predictions.
- **3D Visualization**. The new graphical interface of MutaFrame is designed to facilitate the interpretation of the variant predictions, and is intended for users who do not necessarily have a background in structural bioinformatics. Of particular interest is the visualization of the variants in the context of the protein three-dimensional (3D) structures, which allows users to immediately identify the interactions between the mutated amino acids and the neighboring residues and their position in the structure.

More details on these three utilities are given in the next section. Moreover, we showcase in the last section the application of MutaFrame to the Niemann-Pick disease (NPD) through the analysis of variants in the SMPD1 gene that codes for sphingomyelin phoshodiesterase 1. This protein is crucial in lipid metabolism and its disruption causes a vast array of symptoms, ranging from hepatosplenomegaly to mental retardation and infantile death. We show how MutaFrame can help interpret the impact of the variants on the molecular phenotypes and on the disease mechanisms. Note that a predictor of Niemann-Pick disease severity that uses both SNPMuSiC and DEOGEN2 has recently been developed [8].

### MutaFrame Structure

User queries in MutaFrame start from a protein sequence or a UniProt [9] accession number, and a variant in the protein sequence. Upon submission, MutaFrame automatically maps the variant position to the corresponding gene and to the protein structure (if available), collects gene and protein characteristics and annotations, and runs all utilities on the basis of sequence, structure and contextual features.

MutaFrame first provides three main sections describing the effect of the query variant:

- *General information* gives insight into the physicochemical characteristics of the wild-type and mutated residues. It showcases a similarity index between these two residues, as illustrated in Fig. 2.a. Clicking on the similarity bar unravels it into its components, including the BLOSUM62 matrix element [10], as well as the difference in residue size, electric charge and hydrophobicity.
- *DEOGEN* contains the results related to DEOGEN2. The first output is the predicted deleteriousness score, between 0 and 1. Scores higher than 0.5 indicate deleterious variants and scores lower than this threshold, neutral variants. Again, clicking on the bar unravels into its components, which are listed in Table 1. More information is given in the Interpretation Utility section.
- *SNPMuSiC* reports the deleteriousness score of the target variant obtained by the SNPMuSiC predictor, if the variant is introduced in a region of the protein with experimental or modeled structure, as explained in the Interpretation Utility section. Positive and negative values indicate deleterious and neutral predictions, respectively. The solvent accessibility (in %) of the wild type residue is also reported.

**Figure 2:**
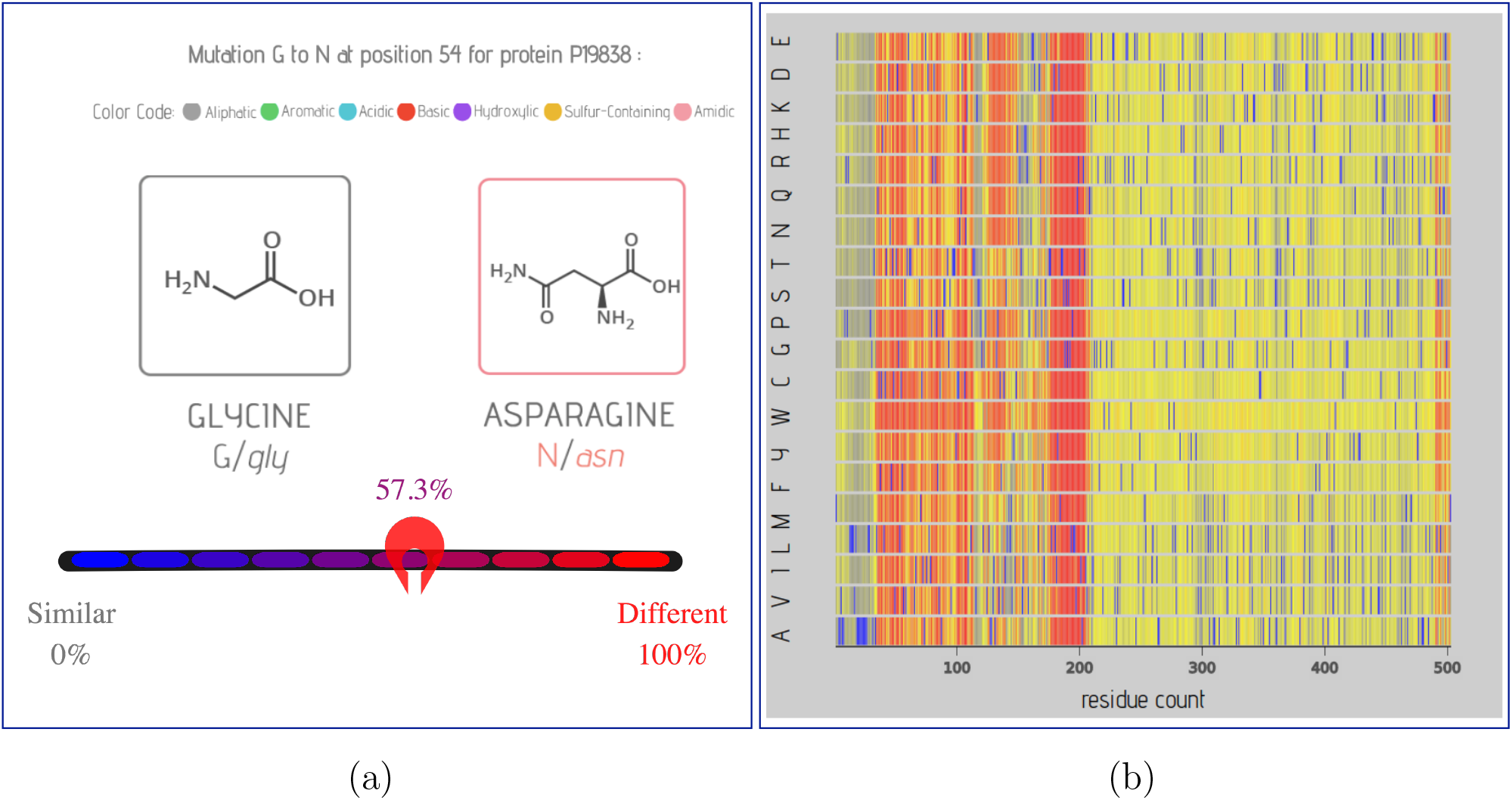
Visualization of MutaFrame’s results. (a) General information about wild-type and variant residues; (b) DEOGEN2 score for all possible variants as a function of the position in the sequence, where red means deleterious and blue benign.

**Table 1:** List of features used by DEOGEN2 to predict the deleterious impact of variants. The sequence-based information is in blue and the contextual information in green. MSA means Multiple Sequence Alignments, and PFAM Protein Families.

| Features | Based on | Type | Ref. |
| --- | --- | --- | --- |
| PRovean algorithm [PR] | Amino acid conservation in MSA of homologous proteins | Sequence<br>Per residue | [11] |
| Conservation Index [CI] | Measure of sequence variability in MSA of homologous proteins | Sequence<br>Per residue | [6, 12] |
| mutated/wild-type Log-Odd ratio [LO] | Log-odd ratio of mutated/wild-type residue at given position in MSA | Sequence<br>Per residue | [11] |
| Early Folding [EF] | Predicted mutational impact on the protein folding process | Sequence<br>Per residue | [13] |
| Residual Variation intolerance [RV] | Gene ranking based on tolerance to functional genetic variation | Sequence<br>Per gene | [14] |
| PFam score [PF] | Log-odd ratio of frequency of neutral/deleterious variants in PFAM domains | Context<br>Per domain | [15] |
| INteraction patches [IN] | Annotations of residues belonging to protein-protein interaction patches | Context<br>Per residue | [16] |
| Gene Damage index [GD] | Cumulative mutational gene damage in the general population | Context<br>Per gene | [17] |
| REcessiveness index [RE] | Predicted degree of tolerance to loss of gene function | Context<br>Per gene | [18] |
| gene ESsentiality [ES] | Degree of essentiality from knock-out experiments in mice | Context<br>Per gene | [19] |
| PAthway score [PA] | Log-odd ratio of pathway sensitivity to deleterious variants | Context<br>Per pathway | [6, 12] |

MutaFrame does not only give DEOGEN2 and SNPMuSiC predictions on the target variant, but also average per-residue predictions on all possible variants in the target sequence, which is very useful to obtain a global view of the susceptibility of the protein to deleterious variants. We describe in detail below the different utilities of MutaFrame.

All the MutaFrame results are easily downloadable for offline analyses. Clicking on the gear next to a figure opens a new panel which allows the user to download the figure and the data used to construct it. The “json” and “csv” buttons download the data in JSON and CSV formats, respectively, and the “camera” button downloads the figure in SVG format.

### 1. 3D Visualisation Utility

The MutaFrame webserver has a user-friendly web interface which facilitates decryption and interpretation of the results. A very useful utility is the visualisation of protein 3D structures and the localization of target variants, as illustrated in Fig. 3, since this greatly helps contextualising the amino acid within the protein, and the molecular effects of the variant. We describe below in more detail how the structures of target sequences are retrieved and the main associated tools.

**Figure 3:**
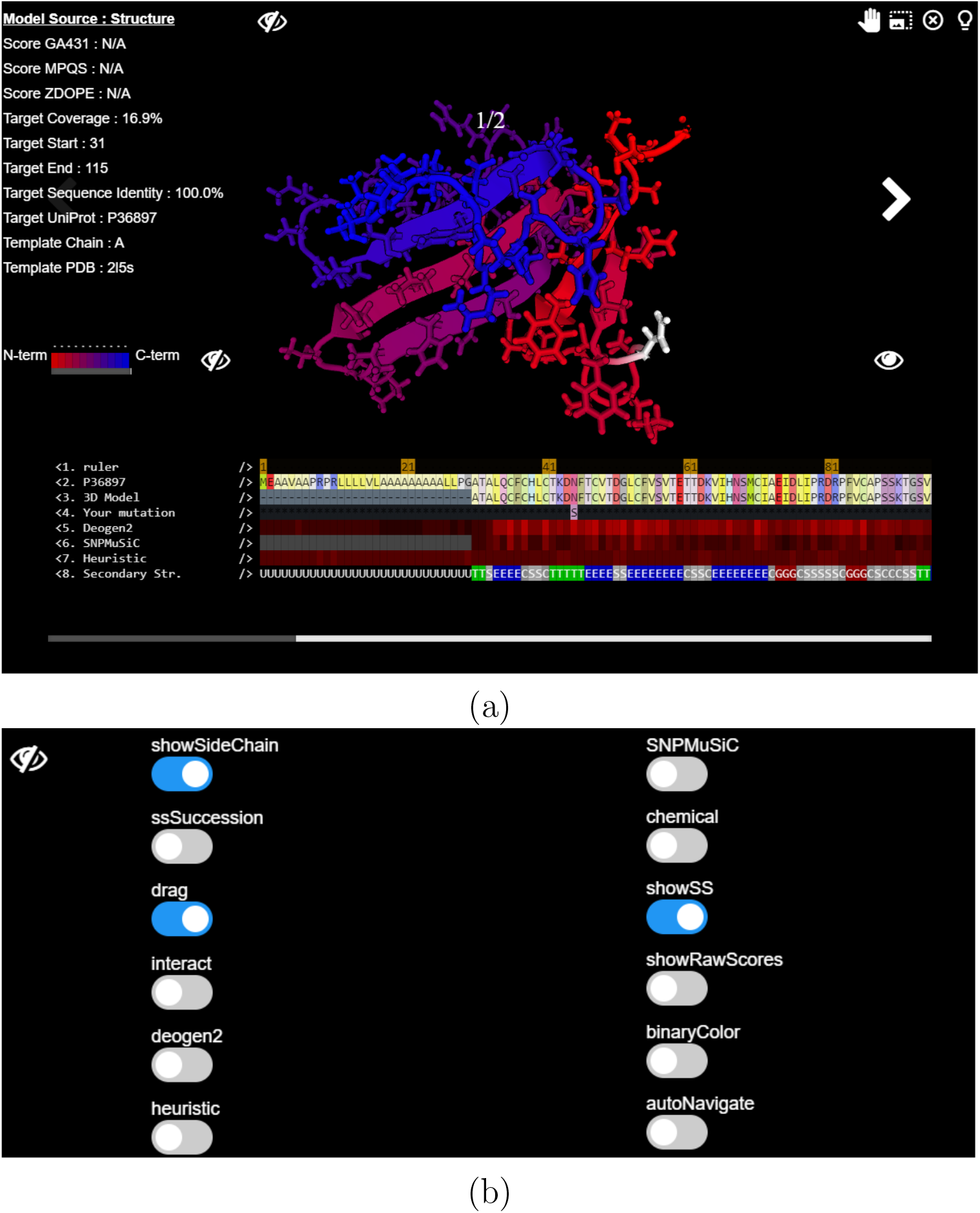
Protein structure visualizer. The “eye” buttons are used to display or hide the different panels. (a) Experimental or modeled 3D structure and sequence alignement between UniProt and PDB sequences; (b) Option panel. The significance of the buttons is described in Table 2.

#### Protein 3D structures of the human proteome

We retrieved structural information of the full human proteome from the Interactome3D dataset [20]. This dataset contains a set of experimentally resolved protein structures as well as homology-modeled structures for which a structural template with a sufficient level of sequence identity and coverage with respect to the target sequence was found in the PDB. When several models were found in Interactome3D for the same protein region, we selected a single representative structure based of Interactome3D’s ranking system that uses a combination of sequence identity and coverage with respect to the template.

As the residue numbers are generally different for protein sequences retrieved from UniProt and from experimental or modeled PDB files, we aligned them using ClustalW [21]. This alignment is shown under the structures (Fig. 3.a).

The final set of structures includes 15,635 entries for the 59,262 UniProt sequences. Out of these 15,635 structures, one half (7,789) are experimental structures and the other half (7,846) are homology models.

#### Structure information

Information about the protein structure is given in the structure visualizer (Fig. 3.a). First, the PDB code and chain name of the experimental structure or of the template used for homology modeling are indicated. In the latter case, the modeling scores GA431 and MPQS obtained from the Interactome3D database are also shown, as well as the start and end residues of the alignment between the target and template sequences, and the sequence identity and coverage.

If several structures exist for the different regions of the target protein, all are shown and it is possible to switch from one to the others without restarting the visualizer. These structures in PDB format and the scripts to color them using PyMol (The PyMOL Molecular Graphics System, Version 1.2r3pre, Schrödinger, LLC) based on the DEOGEN2, SNPMuSiC or heuristic scores can be downloaded by clicking on the download button in the top right corner of the viewer window.

#### 3D structure visualizer

The visualizer in itself shows the protein structure(s) of the target protein, with the wild-type residue at the position of the mutation introduced by the user in white (Fig. 3.a). It offers the possibility to zoom in or out and to rotate the structure. Double clicking on a residue centers the view around it. Hovering over a residue triggers a display mentioning its position and type. By default, the structure is colored using a blue to red gradient from N- to C-terminus.

Different buttons are available to modify the display, listed in Table 2 and shown in Fig. 3.b. They allow showing the secondary structures, the residue side chains, and coloring the protein chain according to the mean per-residue DEOGEN2 or SNPMuSiC score, or according to the mean similarity heuristics. These options are illustrated in Fig. 3.b and can be displayed in the visualizer by clicking on the white “eye” on the right.

**Table 2:** List of options for the MutaFrame protein structure visualizer.

| Option Name | Description |
| --- | --- |
| showSideChain | Toggle display of the side chains of all residues in the protein structure |
| showSS | Display secondary structures as cartoon |
| ssSuccession | Color successions of secondary structures (if showSS enabled) |
| drag | Alignment can be moved around by if enabled |
| interact | Enables display of residue type (when hovering over structure) and per-residue scores (when clicking on prediction rows in alignment) |
| chemical | Color the structure according to the physicochemical properties of the residues |
| heuristic | Color the structure according to the mean residue similarity index |
| deogen2 | Color the structure according to the mean DEOGEN2 score per residue |
| snpMusic | Color the structure according to the mean SNPMuSiC score per residue |
| showRawScores | Enables display of raw scores when clicking on the prediction rows (in alignment) |
| binaryColor | Change color of scores in the alignments: red/green or shades of red |
| autoNavigate | Move the alignment when hovering over a residue if enabled (if interact enabled) |

#### Sequence alignments

The alignment of the target protein sequence with the subset of residues that are part of the protein structure(s) are shown below the 3D structures, as illustrated in Fig. 2.a. Underneath the sequence alignment, the mean per-residue scores of the DEOGEN2 and SNPMuSiC predictors and of the heuristic similarity are given, using a light to dark red scale. This color code can be changed using the *binary-Color* option, making neutral mean scores green and deleterious ones red (Table 2). Clicking on one of the colored blocks displays the average per-residue prediction (deleterious or neutral) for the residue. The *showRawScores* option can be used to display the average per-residue scores (numeric values).

### 2. Interpretation Utility

The core of the MutaFrame webserver provides accurate predictions of the deleterious or neutral nature of any missense variant in any protein of the human exome. Two complementary predictors, DEOGEN2 [5] and SNPMuSiC [7], fulfill this role. The features used by these predictors are of three categories: sequence-, structure- and context-based. We briefly review how these predictors work, the features they exploit, and the information they provide on the impact of variants at the molecular and phenotypic levels.

- **DEOGEN2** is a deleteriousness variant predictor which requires the protein sequence as input. It is based on the set of sequence and contextual features listed in Table 1 and a random forest classifier with 200 trees to construct a prediction model. The MutaFrame webserver returns not only the overall prediction of DEOGEN2, but also a careful analysis of the contribution of each of the features to the global score, as illustrated in Fig. 1.b. In this way, it is possible to rationalize the results and have an explanation about the reasons behind the deleteriousness or neutrality of the variant. For example, if one of the features such the protein-protein interaction patch feature (IN, see Table 1) drives the predictions, it is likely that the variant impacts on the interaction of its host protein with one of more other proteins. More precisely, the main DEOGEN section contains the following results:
  1. A barplot displaying the contribution of each feature to the global DEOGEN2 score (Fig. 4.a). These values were obtained by an automated interpretation of the random forest model used by DEOGEN2; see [5] for details.
  2. A barplot displaying the raw values of all features (Fig. 4.b). These are the values used as input by DEOGEN2.
  3. A histogram that displays the number of variants in the target protein that have a specific deleteriousness score.
  4. A heatmap representation of the scores of all possible variants along the protein sequence (Fig. 2.b). This representation can switch between mean scores per position or individual variant scores by clicking on the “mode” button on the graph. Note that the graphs are interactive and that additional information is displayed by hovering the cursor over the corresponding bar. The DEOGEN2 performances are described in a section below. More details about the model, its characteristics and applications can be found in [5].
- **SNPMuSiC** [7] is a deleteriousness variant predictor that requires an experimental or modeled 3D structure of the target protein as input. For experimental structures, SNPMuSiC uses the biological unit reported in the PDB, while for modeled structures, it uses the modeled chain only. SNPMuSiC is based on both structural and evolutionary information and outputs a stability-driven deleteriousness index of the query variant.

**Figure 4:**
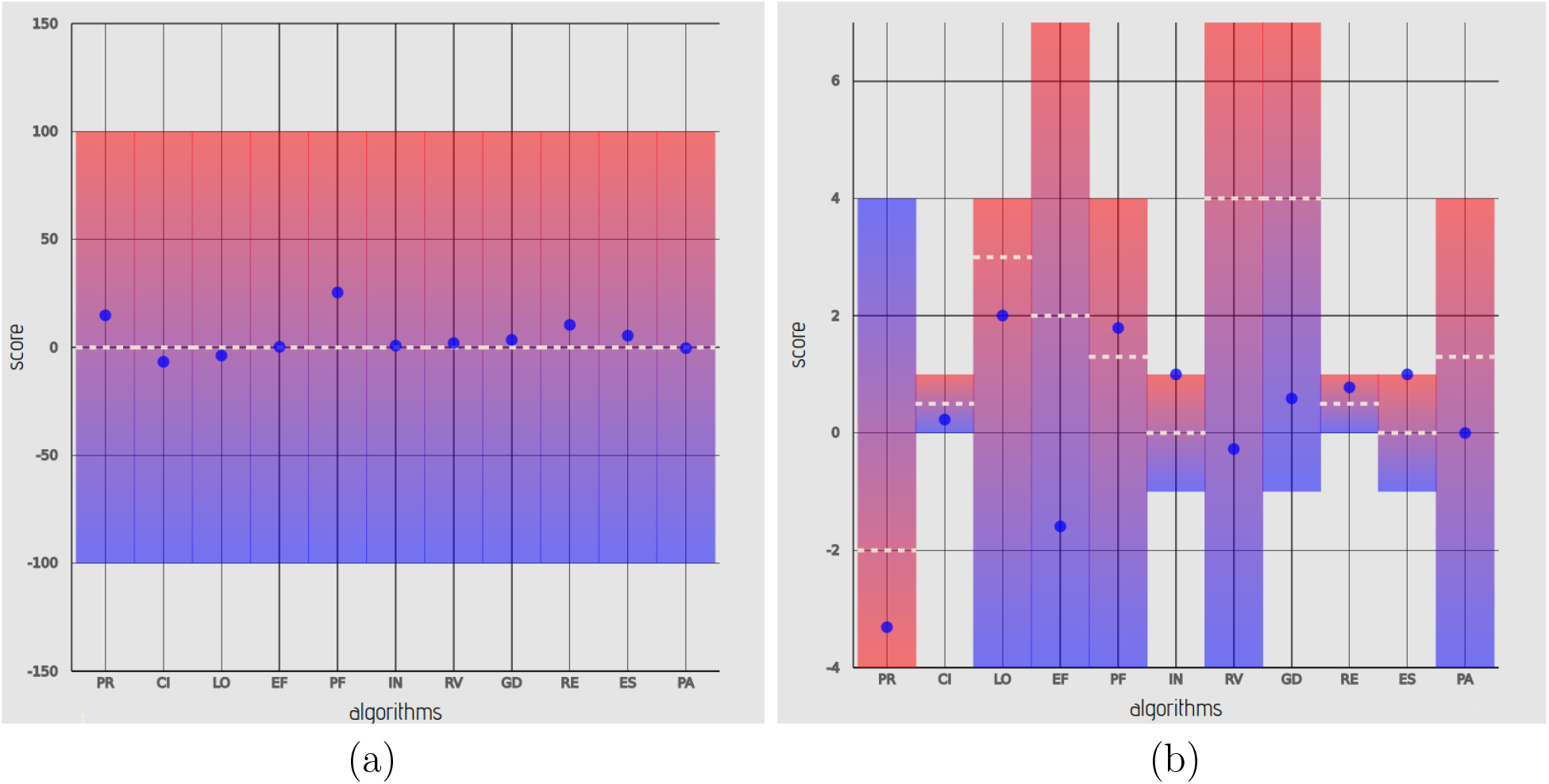
Feature values and their contributions to DEOGEN2 predictions. The results were obtained for the N45S variant in the P36897 protein. (a) Individual contribution of each feature (listed in Table 1) to the final DEOGEN2 prediction; (a) raw values of the features.

To perform its predictions, SNPMuSiC uses a series of statistical potentials, which are coarse-grained mean force potentials widely applied in protein science [22]. They are computed from frequencies of associations between a sequence feature (type of amino acid(s)) and a structural feature (main chain torsion angle domain(s), residue solvent accessibility(ies), spatial distance(s) between residue pairs) in a dataset of experimental 3D protein structures, which are turned into folding free energies using the Boltzmann law (see [23] for technical details).

A series of 13 different potentials are used to estimate the difference in folding free energy between the wild-type and variant residues (noted ΔΔ*W*). These energy values are combined using artificial neural network techniques, and then integrated with the evolutionary score of the PROVEAN algorithm [11] based on residue conservation across natural evolution, to predict the deleterious or neutral nature of the target variant. The list of the features used by SNPMuSiC is showed in Table 3.

**Table 3:** List of all the features used in the SNPMuSiC model to predict variant deleteriousness. In green, the structure-based, stability-driven features. In blue, the sequence-based, evolutionary-driven feature.

| Feature type | Name | Element(s) |
| --- | --- | --- |
| Torsion angle potentials | $\Delta W_{st}$<br>$\Delta W_{stt'}$<br>$\Delta W_{ss't}$ | $t, t' =$ backbone torsion angle domain<br>$s, s' =$ amino acid type |
| Solvent accessibility potentials | $\Delta W_{sa}$<br>$\Delta W_{saa'}$<br>$\Delta W_{ss'a}$ | $a, a' =$ residue solvent accessibility<br>$s, s' =$ amino acid type |
| Torsion angle/<br>solvent accessibility potentials | $\Delta W_{sta}$ | $a =$ residue solvent accessibility<br>$t =$ backbone torsion angle domain<br>$s =$ amino acid type |
| Distance potentials | $\Delta W_{sd}$<br>$\Delta W_{sds'}$ | $d =$ distance between residue pair<br>$s, s' =$ amino acid type |
| Distance/<br>torsion angle/<br>solvent accessibility potentials | $\Delta W_{sad}$<br>$\Delta W_{std}$<br>$\Delta W_{sads'a'}$<br>$\Delta W_{stds't'}$ | $d =$ distance between residue pair<br>$a, a' =$ residue solvent accessibility<br>$t, t' =$ backbone torsion angle domain<br>$s, s' =$ amino acid type |
| Volume terms | $\Delta V_+$<br>$\Delta V_-$ | $V =$ amino acid volume<br>$\Delta V_+(\Delta V_-) = \Delta V$ if $\Delta V \geq 0 (< 0)$ |
| PROVEAN | $PR$ | Amino acid conservation<br>in MSA of homologous proteins |

SNPMuSiC’s predictions are thus essentially based on the impact that a variant has on protein stability. This means that, if it predicts a variant to be deleterious, it is likely that the mutated protein is destabilized and undergoes a modification in its conformation that affects function; or, alternatively, that it is too stabilized and lacks the necessary flexibility and/or the specific functional residues to function properly. As a consequence, a drawback of this predictor is its inability to predict variants that are deleterious due to other factors than stability. As an example, SNPMuSiC is unable to predict a mutation to be deleterious if its molecular effect is to disrupt a protein-protein or protein-ligand interaction necessary for function. However, its contributions perfectly integrate into the MutaFrame framework and provide fundamental information about the variant effects, since stability is well known to be one of the main contributions to folded protein fitness [24].

### 3. Mapping Utility

MutaFrame provides two computational tools to cross reference different sources of information at the variant, gene, protein sequence, 3D structure, and phenotypic levels. They are described below:

- The protein **structure visualizer** shows the experimental or modeled structures of the target protein, if available (Fig. 3.a). This tool displays an alignment between the residues of the target protein sequence and the subset of residues that are part of the protein structure(s). A detailed description of this tool is given in Section 1.
- **VarCraft** is a tool that maps the variant residues in the target protein to biocurated annotations from the Online Mendelian Inheritance in Man database (OMIM) [25]. The tool can be used in two different ways. Clicking on the VarCraft button opens a small window providing the user with two options: starting the VarCraft mapper or labeling mutations (Fig. 5.a).
  1. Labeling mutations. Go to the score distribution or to the heatmap of the DEOGEN2 predictor described in Section 2. Enter one or several variants in the white box of VarCraft shown in Fig. 5.a. When clicking on the purple box “Label Mutations”, these variants appear in the score distribution or in the heatmap with their DEOGEN2 and heuristic scores, as shown in Fig 5.b-c. Before entering other mutations, click on the “Clear Mutations” box.
  2. VarCraft mapper. Clicking on “Start VarCraft” opens a window and starts a series of mapping procedures between variants annotated in the genome Aggregation Database (gnomAD) (using the genome assembly GRCh38) and in ClinVar [26], the substitutions at the nucleic acid and amino acid levels, the positions in the chromosomes, genes and proteins, and the OMIM annotations. Once the process is finished, an interactive graph appears as illustrated in Fig. 6, which shows the position (Pos) of the variant in the chromosome, the wild-type nucleotide (Ref), the mutant nucleotide (Alt), its allele frequency (more precisely, − log(allele frequency)) with missing values (indicated as NaN) meaning that the variant was not found in gnomAD (pAF), the cDNA variant identifier (cDNA), the protein variant in Human Genome Variation Society (HGVS) format [27] in the gnomAD transcript (pHGVS) and in the amino acid sequence considered in the MutaFrame database (MtfPos), the OMIM annotation when available (OMIM), the variant identifier in ClinVar [26] when available (ClinVar) and the DEOGEN2 score (DEOGEN2). By hovering the mouse over a particular line, the tool zooms on this line and provides the numerical values (Fig. 6b). By clicking on the gene symbol, the user can retrieve the Uniprot file used in MutaFrame database. The gene symbol, gene identifier, OMIM identifier and transcript identifier are linked to the entry in Ensembl, OMIM and gnomAD, respectively. All the data shown in VarCraft can be downloaded by clicking on the download button in the top right corner of the VarCraft window.

**Figure 5:**
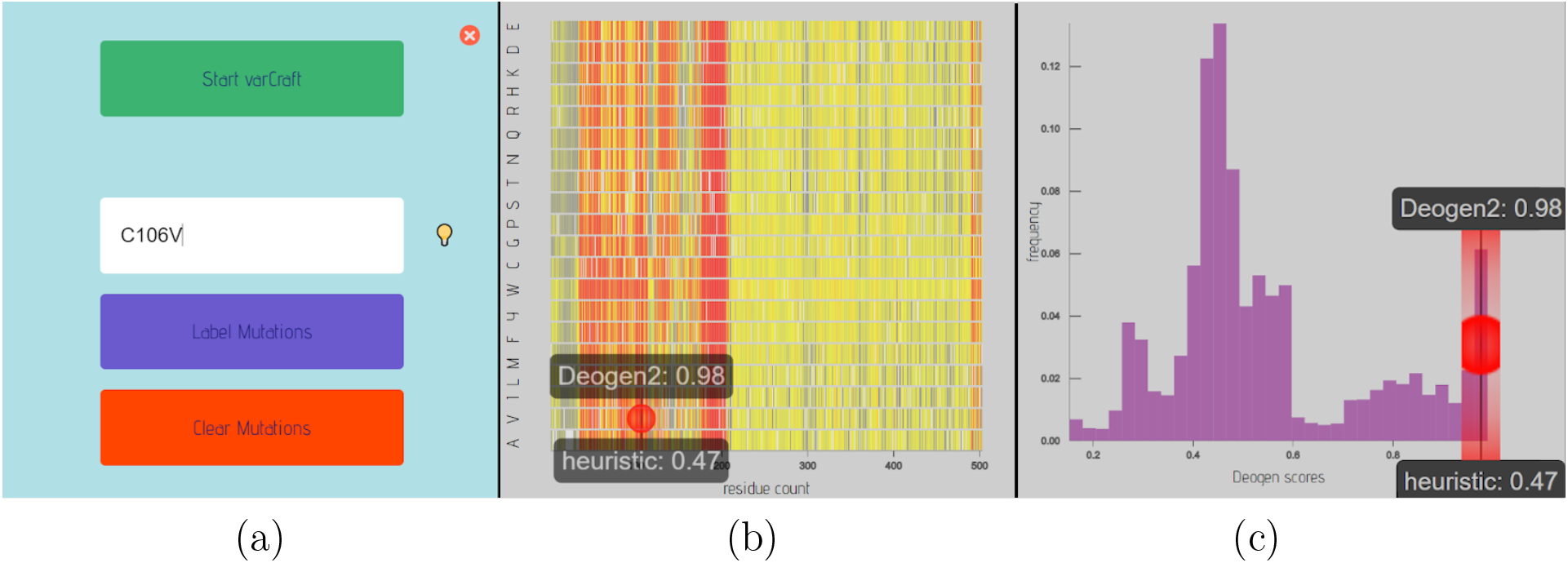
(a) Window appearing when clicking on the VarCraft button. It is possible to label mutations in the DEOGEN2 heatmap (b) or in the DEOGEN2 scores distribution (c) using the “Label Mutations” button. The “Clear Mutations” button removes the label from the graphs. Here, the variant C106V of Uniprot P36897 was labelled.

**Figure 6:**
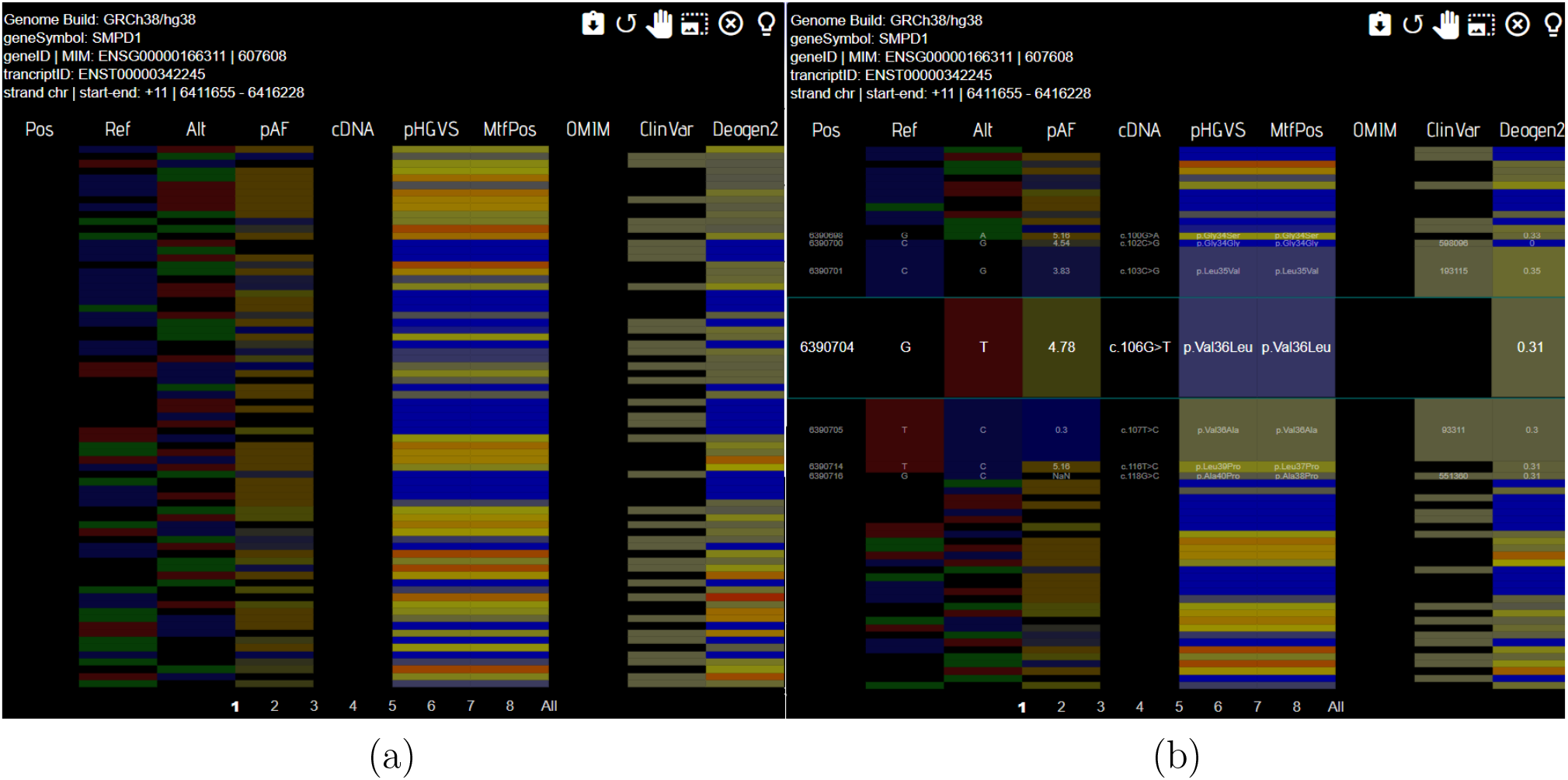
VARCRAFT interactive visualization of annotations. (a) Annotations for the SMPD1 gene. “Pos” column: position number in the chromosome; “Ref”: reference nucleotide (green = “A”, black = “G”, blue = “C”, red = “T”); “Alt”: variant nucleotide (same color code); “pAF”: -log(Allele Frequency) (blue/yellow scale with blue meaning high allele frequency and low pAF; NaN means that the variant is not in gnomAD and thus that we do not have access to its allele frequency; “cDNA” variant identifier in the cDNA sequence; “pHGVS” and “MtfPos”: variant identifier at protein level in HGVS nomenclature [27] in the gnomAD transcript and in the transcript considered in MutaFrame, respectively (blue/red scale depending on similarity between reference and variant residues, with blue meaning high similarity); “OMIM”: annotation identifier in the OMIM database; “ClinVar”: variant identifier in the ClinVar database (color showing the presence or absence of an identifier); “Deogen2”: DEOGEN2 score (blue/red scale depending on the DEOGEN2 score, where blue is neutral and red deleterious). (b) Hovering the cursor over a variant unfolds the row to show more details.

## DEOGEN2 and SNPMuSiC performances

An extensive evaluation of the performances of both deleteriousness predictors DEOGEN2 and SNPMuSiC has been presented in [5] and [7], respectively. Here we compared the predictions of these two methods to assess their strengths, weaknesses and complementarity. We did this analysis on the basis of the dataset constructed in [7], which contains 5,192 human variants for which the predictions of both predictors are available.

The overall performances of SNPMuSiC and especially DEOGEN2 are good, as shown in Table 4. The balanced accuracy (BACC) is equal to 77% and 89% and the positive predicted value (PPV) is above 90% for both predictors. Note that these values are in direct validation, but that cross-validated scores are only 2% lower at most [5, 7]. In the subset of variants for which the two predictors agree, which contains 4,123 of the 5,192 variants, the scores significantly improve: the BACC reaches 92% and the PPV 97%.

**Table 4:** Balanced Accuracy (BACC) and Positive-Predictive Value (PPV) of DEOGEN2 and SNPMuSiC, computed on SNPMuSiC’s learning dataset [7]. Consensus is computed on the subset of variants for which the predictions of DEOGEN2 and SNPMuSiC both agree.

| Predictor | BACC | PPV |
| --- | --- | --- |
| SNPMuSiC | 77% | 90% |
| DEOGEN2 | 89% | 95% |
| Consensus | 92% | 97% |

The confusion matrices for SNPMuSiC, DEOGEN2 and the consensus predictions are reported in Tables 5, 6. and 7. The lesser performances of SNPMuSiC are likely related to its ability to only predict variants of which the deleteriousness is caused by stability issues of the 3D structure, as previously discussed [7]. It is thus expected that SNPMuSiC overpredicts neutral mutations.

**Table 5:** SNPMuSiC’s confusion matrix, computed on SNPMuSiC’s learning dataset [7].

| SNPMuSiC | Annotation |  |  |
| --- | --- | --- | --- |
| Predicted | Disease | Neutral | Total |
| Disease | 3122 | 353 | 3475 |
| Neutral | 769 | 948 | 1717 |
| Total | 3891 | 1301 | 5192 |

**Table 6:**
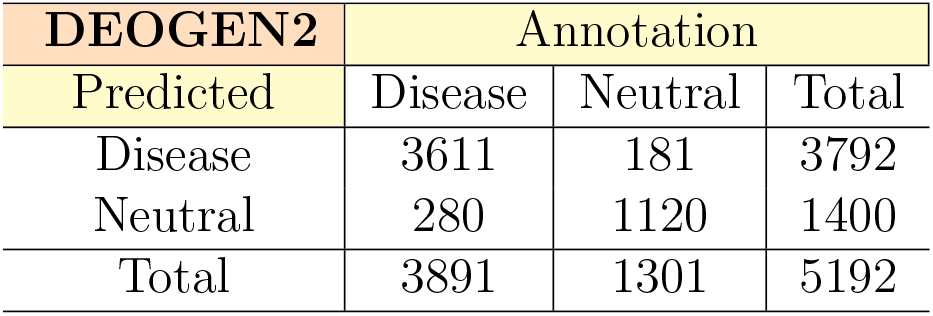
DEOGEN2’s confusion matrix, computed on SNPMuSiC’s learning dataset [7].

**Table 7:**
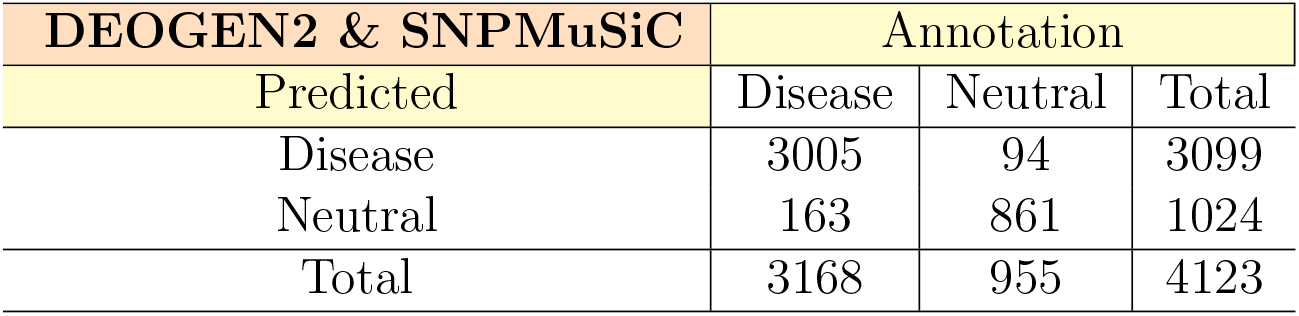
DEOGEN2 and SNPMuSiC confusion matrix for consensus predictions, computed on SNPMuSiC’s learning dataset [7].

This effect is clearly visible in Fig. 7, where the distribution of the SNPMuSiC and DEOGEN2 scores are shown separately for neutral and deleterious mutations. Indeed, for the whole series of deleterious variants that are correctly predicted by DEOGEN2, but not by SNPMuSiC, we can assume they are likely deleterious for other reasons than stability. There are also some mutations that are correctly predicted by SNPMuSiC and not by DEOGEN2, in which stability is expected to play a central role.

**Figure 7:**
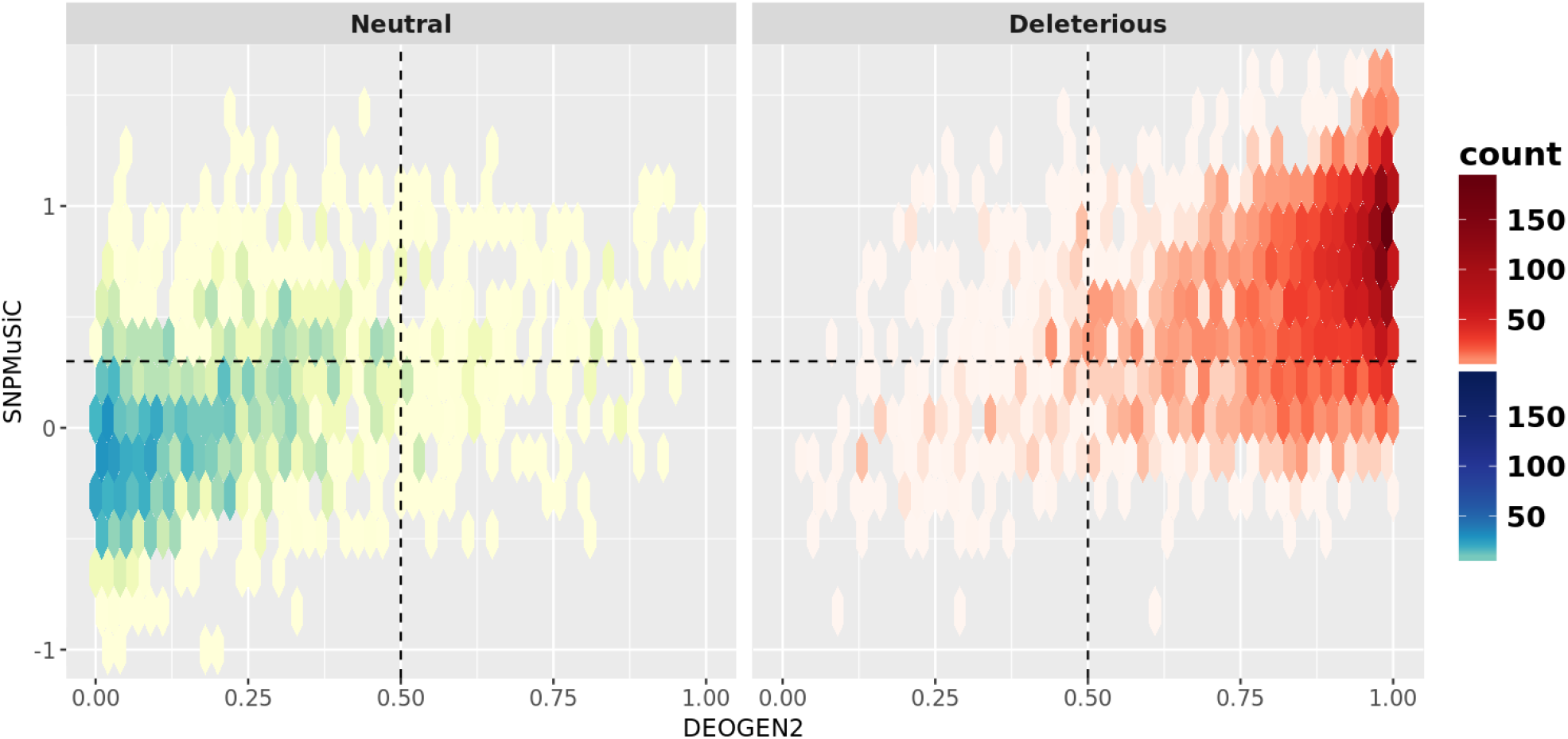
Distribution of DEOGEN2 scores (x-axis) and SNPMuSiC scores (y-axis) scores for deleterious variants (in red) and neutral variants (in blue), computed on SNPMuSiC’s learning dataset [7]. Dotted lines represent the prediction thresholds of the predictors.

In conclusion, there are several advantages to the combination of the two predictors: (1) when there is a consensus between the predictors, the prediction reliability is very high; (2) when SNPMuSiC predicts a variant as deleterious, we can interpret its molecular effect to be a significant change in stability; (3) the predictions of DEOGEN2 can be unraveled in terms of contextual or evolutionary features. We thus get an improved explanatory power due to the combination of the two predictors that integrate structure-based information from SNPMuSiC and evolutionary and contextual information from DEOGEN2.

## Case study

In this section, we illustrate the application of the MutaFrame webserver to variants in lysosomal acid sphingomyelinase, also known as sphingomyelin phosphodiesterase (SMPD1, UniProt id P17405). Variants in SMPD1 are known to cause the Niemann-Pick disease (NPD) of types A and B, which are lipid storage disorders characterized by various clinical phenotypes such as hepatosplenomegaly and pulmonary insufficiency [28]. A computational variant analysis related to NPD severity can be found in [8].

More precisely, we show here how to use MutaFrame to investigate the impact of the two variants Arg228Cys and Trp244Cys in SMPD1.

- **Arg228Cys** is a known deleterious mutation [29, 30, 31] reported in gnomAD and ClinVar and its molecular effect results in a residual activity of about 5% with respect to the wild type protein [29]. Submitting this variant to the MutaFrame webserver yields the consensus prediction that it is deleterious. Indeed, DEOGEN2 classifies the variant as highly deleterious with a score of 0.906 (with respect to a threshold value of 0.5) and SNPMuSiC classifies it as strongly deleterious with a score 0.64 (with respect to a threshold value of 0). To gain insight into the molecular effect of the variant, we analyzed further information provided by MutaFrame. From the DEOGEN2 feature analysis, we learned that the prediction is dominated by evolutionary terms, especially the PROVEAN score and the mutated/wild-type log-odd ratio. This suggests that the wild-type Arg has a functional or structural role. In addition, SNPMuSiC predicts the Arg228Cys variant protein to be substantially less stable than the wild-type. This suggests an impact of this variant on protein stability. Using the 3D visualization tool, we can furthermore zoom on the spatial region surrounding Arg228 as illustrated in Fig. 8. We can clearly see that this Arg interacts with Asp210, forming a salt bridge, which is broken upon mutation. These different results thus point towards structure destabilisaton upon mutation.
- **Trp244Cys** is another known deleterious mutation reported in UniProt (but not in gnomAD and ClinVar, which explains that it is absent from VarCraft), which reduces the relative enzymatic activity to less than 10% [32, 33]. Again, both predictors predict the Trp244Cys variant as strongly deleterious, DEOGEN2 with a score of 0.881 and SNPMuSiC with a score of 1.06. The evolutionary features of DEOGEN2 indicate that Trp244 is strongly conserved, and SNPMuSiC predicts that the Trp244Cys mutation has an important impact on protein stability. The structural analysis using MutaFrame’s visualization tool shows that Trp244 is involved in an interaction network of aromatic and positively charged residues, as illustrated in Fig. 8. Indeed, Trp244 forms a cation-*π* interaction with Arg255 and a T-shaped *π*-*π* interaction with Tyr243. Breaking these interactions upon Trp244Cys mutation has an impact on the ability of SMPD1 to maintain its conformation and enzymatic function, even though the mutation is relatively far from the functional site.

**Figure 8:**
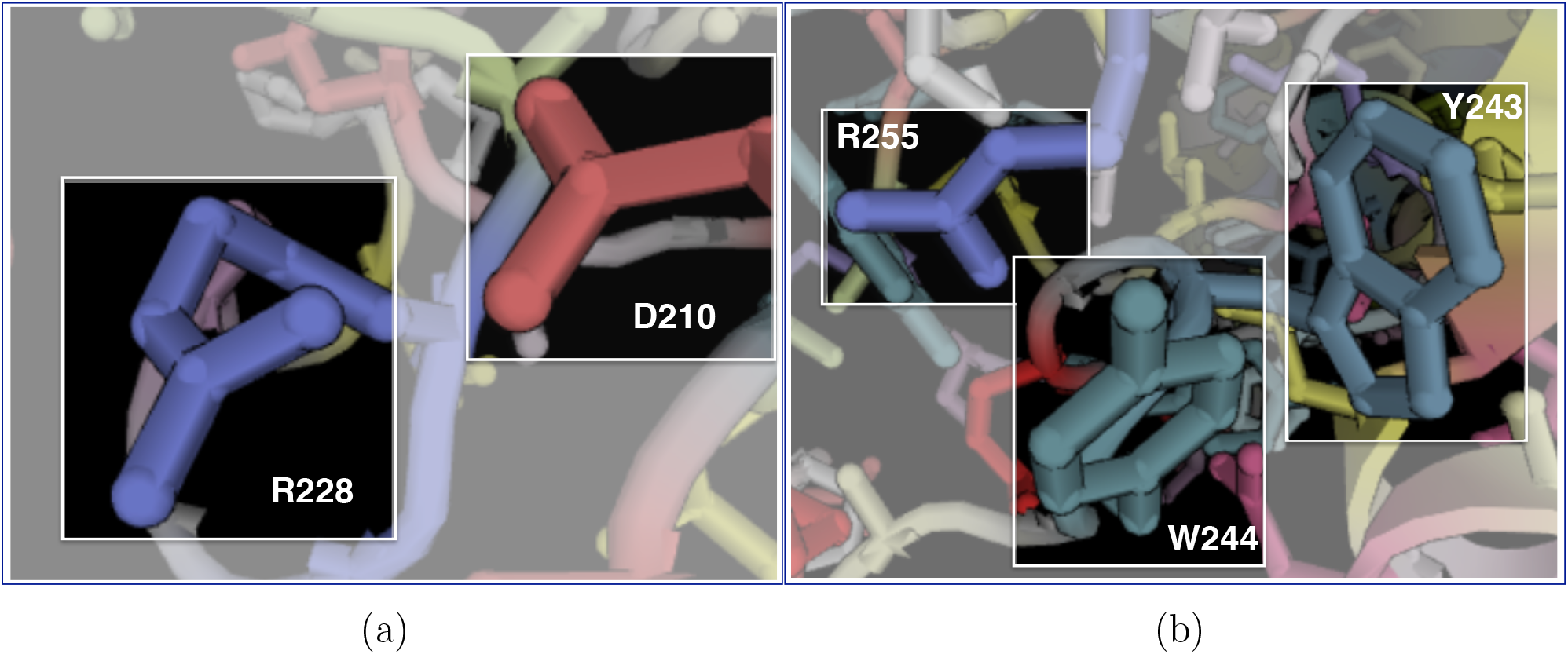
Sphingomyelin phosphodiesterase (SMPD1) involved in the Niemann-Pick disease (PDB code 5JG8). (a) Environment of Arg 228, which interacts with Asp 210 through a salt bridge. (b) Environment of Trp 244, which forms a cation-*π* interaction with Arg 255 and a *π*-*π* interaction with Tyr 243.

